# Probabilistic Associative Learning Suffices For Learning The Temporal Structure Of Multiple Sequences

**DOI:** 10.1101/545871

**Authors:** Ramon H. Martinez, Anders Lansner, Pawel Herman

**Affiliations:** KTH Royal Institute of Technology Computational Brain Science Lab Lindstedtsvägen 5, 10044 Stockholm, Sweden; Stockholm University, Mathematics and KTH Royal Institute of Technology, Computational Brain Science Lab Lindstedtsvägen 5, 10044 Stockholm, Sweden

**Keywords:** Keywords neural networks, associative learning, sequence learning, attractor models

## Abstract

Many brain phenomena both at the cognitive and behavior level exhibit remarkable sequential characteristics. While the mechanisms behind the sequential nature of the underlying brain activity are likely multifarious and multi-scale, in this work we attempt to characterize to what degree some of this properties can be explained as a consequence of simple associative learning. To this end, we employ a parsimonious firing-rate attractor network equipped with the Hebbian-like Bayesian Confidence Propagating Neural Network (BCPNN) learning rule relying on synaptic traces with asymmetric temporal characteristics. The proposed network model is able to encode and reproduce temporal aspects of the input, and offers internal control of the recall dynamics by gain modulation. We provide an analytical characterisation of the relationship between the structure of the weight matrix, the dynamical network parameters and the temporal aspects of sequence recall. We also present a computational study of the performance of the system under the effects of noise for an extensive region of the parameter space. Finally, we show how the inclusion of modularity in our network structure facilitates the learning and recall of multiple overlapping sequences even in a noisy regime.

## 1 Introduction

From throwing spears in the savanna to the performance of a well rehearsed dance, human behavior reflects an intrinsic sequential structure. In this light, is not surprising that sequential activity has been found in the neural dynamics across different anatomical brain areas such as the cortex (Luczak et al., 2007; Jin et al., 2009; Harvey et al., 2012; Tang et al., 2008), the basal ganglia (Barnes et al., 2005; Mello et al., 2015; Gouvêa et al., 2015; Bakhurin et al., 2017; Dhawale et al., 2017; Rueda-Orozco and Robbe, 2015; Jin et al., 2009), the hippocampus (Nádasdy et al., 1999; Pastalkova et al., 2008; Louie and Wilson, 2001; Davidson et al., 2009; MacDonald et al., 2013) and the HVC area in songbirds (Hahnloser et al., 2002; Kozhevnikov and Fee, 2007). Moreover, sequential activity is not only present in a wide range of neuroanatomical areas but is also associated with an ample repertoire of behaviors and cognitive processes including sensory perception (Jones et al., 2007; Crowe et al., 2010), memory (Abeles et al., 1995; Seidemann et al., 1996; Fujisawa et al., 2008), motor behavior (Averbeck et al., 2002; Nakajima et al., 2009) and decision making (Lapish et al., 2008; Harvey et al., 2012). In our view, the entanglement of sequential activity with cognitive processes and behavior strongly suggests that sequential activity is an essential component of the information processing capabilities of the brain and therefore demands better understanding. A plausible hypothesis for the ubiquity of sequential activity is a common learning mechanism for the construction of temporal representations at the network level. Inspired by experimental evidence we propose the following constraints and properties for the neural representations and the underlying network mechanisms: First, the recall dynamics of a sequence should reflect key temporal features of the input or training signal (Johnson et al., 2010). Second, the network should enable temporal scaling, that is, once a sequential representation has been learned, internal neural network’s mechanisms should suffice to contract or dilate its recall duration (Euston et al., 2007; Ji and Wilson, 2007). Finally, as the same neural network circuits have been observed to exhibit many sequential trajectories accounting for different behaviors (Pastalkova et al., 2008), it is desirable for the network to posses mechanisms to store and recall multiple and, to some extent, overlapping sequences (Agster et al., 2002).

There is evidence that sequential activity can be characterized as a succession of meta-stable cell assemblies in the cortex (Seidemann et al., 1996). Attractor neural networks have a long standing tradition as models of sequential activity with meta-stable states corresponding to attractor patterns (Amari, 1972; Willwacher, 1982). Hopfield in his seminal work (Hopfield, 1982, 1984) already noted that an asymmetric connectivity in a recurrent attractor network was conducive to sequential recall. However, in the most basic implementation, the asynchronous update dynamics of these Hopfield models resulted in mixed patterns, thereby gradually diluting sequential recall with time (Kühn and van Hemmen, 1991). To overcome such limitations, temporal traces of the activity were utilized successfully as a mechanism to keep the meta-stable states active for long enough to ensure a successful transition between the patterns and some models even allow for temporal rescaling of the dynamics (Kleinfeld, 1986; Sompolinsky and Kanter, 1986). However, such models are unable to properly integrate the temporal structure of the input due to the discrete nature of their learning rule. A more sophisticated approach relies on systematically considering all the possible delays of the input and calculate all the resulting cross-correlations (Herz et al., 1989; Coolen and Gielen, 1988). While in principle these models are able to lean arbitrary variations in the temporal structure of the input, in practice they are limited by an explosion in the number of parameters as the connectivity matrix scales with the size of the longest transition. In this work we propose an attractor model that uses the following properties to overcome the aforementioned problems: 1) It exploits temporal traces for learning in a probabilistic framework (Tully et al., 2016). The temporal nature of the traces allows us to capture the temporal structure of the input, while avoiding an explosion in the number of parameters by collapsing the temporal structure into statistical estimates of the connectivity. 2) The sequence transition mechanism rests on the meta-stability of the attractor dynamics by means of intrinsic adaptation of the network units coupled with a competition mechanism that bias the transition in the correct direction. At the same time the intrinsic adaptation allows for the internal control and rescaling of the recall dynamics. 3) The use of a modular structure in our network facilitates both flexible learning and recall of overlapping representations.

Several network models have been proposed to account for sequential activity. While Veliz-Cuba et al. (2015) reported that their network could learn the temporal structure of the input, it required a fine-tuned relationship between synaptic, dynamic and homeostatic parameters. Additionally, their model lacked a mechanism for temporal rescaling and the question of learning multiple sequences was not addressed. In a recent approach by Pereira and Brunel (2018) persistent or sequential activity dynamics could be learned depending on the temporal structure of the input. However, the proposed network did not solve the problem of temporal scaling nor the acquisition of multiple sequences. Using spike-time-dependent plasticity (STDP) with heterosynaptic competition Fiete et al. (2010) demonstrated the capability of their model to learn multiple sequences from random activity but handling input with specific temporal structure was not elaborated in their work. Furthermore, Byrnes et al. (2011) addressed the problem of learning overlapping sequences but their approach did not scale well as it relied on a single unique representation for every sequence even if they had overlapping elements. Finally, Murray et al. (2017) proposed an inhibitory network inspired by the basal ganglia that achieves temporal rescaling by means of the interplay between synaptic fatigue and external input. In this model, however, the problem of handling multiple sequences could be solved only by assuming the existence of such representations in an upstream network, which we consider as a strongly limiting factor.

Inspired by our previous modelling efforts to study sequence (Tully et al., 2016) and word list learning phenomena (Lansner et al., 2013) we propose here a modular attractor memory neural network model that learns sequential representations by means of the combination of the Bayesian Confidence Propagating Neural Network (BCPNN) learning mechanism Lansner and Ekeberg (1989) and asymmetrical temporal synaptic traces. We proceed by first presenting the network and its dynamics. Then, we derive analytical formulae for the temporal structure of the recall process in noiseless conditions. We also describe how learning is accomplished in the network through the use of synaptic traces and study how the temporal structure of the input is accounted for in the recall dynamics by means of the BCPNN learning rule. We follow up with a systematic characterization of the effects of noise on the sequence recall capability of the network. Finally, we elaborate on how the modularity of the network enables learning overlapping sequences and discuss key limitations.

## 2 Results

### 2.1 Sequence recall

Following previous work on cortical attractor memory modelling (Tully et al., 2016; Lansner et al., 2013) we present here a network capable of learning, recalling and processing sequential activity. We utilize a population model of the cortex where units represent aggregations of neurons (cortical columns). Consistently with the mesoscale neuroanatomical organization, those units are organized into hypercolumns, where winner-takes-all (WTA) dynamics keeps the activity within the module normalized (Douglas and Martin, 2004). The topological organization of the model is presented in Fig. 1A. The circuit implements attractor dynamics (Lansner, 2009) that leads the evolution of the network towards temporary or permanent patterns of activity. We refer to these stable or meta-stable states as the stored patterns of the network. The patterns themselves are defined by self-recurrent excitatory connectivity that tends to maintain the pattern in place once activated (represented by *w*_*self*_ in Fig. 1B). The patterns can naturally be thought of as cell assemblies distributed among the hypercolumns in the network. The WTA mechanism renders the activity of the units mutually exclusive within the hypercolumns and therefore ensures sparse activity (Foldiak, 2003). Sequential activation of patterns can be induced by feed-forward excitation (represented by *w*_*next*_ in Fig. 1B) coupled with an adaptation mechanism whose role is to cease current pattern activity thereby counteracting the pattern retention effects of the self-recurrent connectivity.

**Figure 1:**
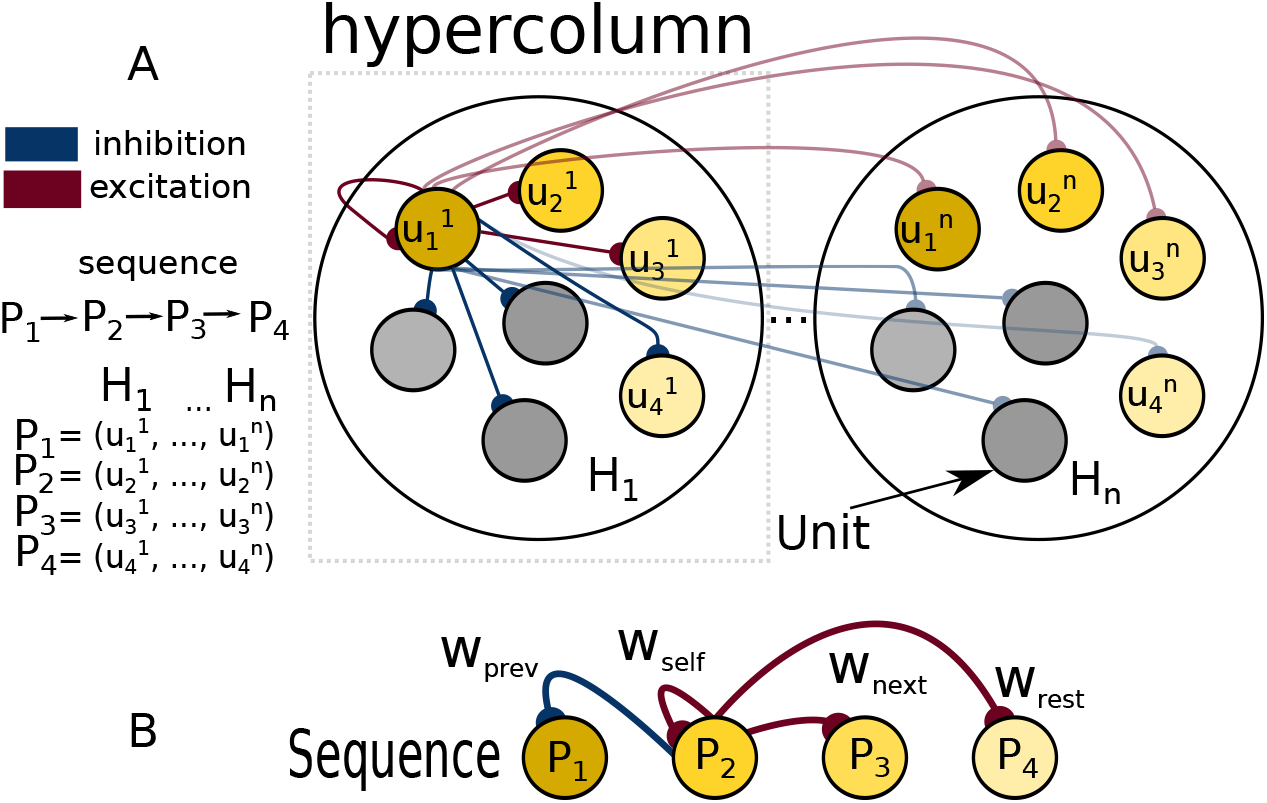
Network architecture and connectivity underlying sequential pattern activation. (A) network topology. Units 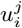 are organized into hypercolumns *H*_1_, *…, H*_*n*_. At each point in time only one unit per hypercolumn is active due to a WTA mechanism. Each memory pattern is formed by a set of recurrently connected units distributed across hypercolumns. For simplicity and without compromising the generality we adopt the convention 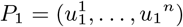. We depict stereotypical network connectivity by showing all the units that emanate from unit 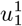. The unit has excitatory projections to the proximate units on the sequence (connections from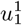 to 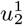 and 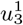 and the corresponding units in other hypercolumns) and inhibitory projections to both the units that are farther ahead on the sequence (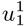 to 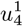) and the units that are not in the sequence at all (gray units). (B) abstract representation of the relevant connectivity for sequence dynamics. Please note that only connections from *P*_2_are shown.

We model the dynamics of the units with a population model equation (Wilson and Cowan, 1972). As described in Eq. 1 the current *s* changes according to the base rate *β*_*j*_ (also called the bias term) plus the total incoming current from the other units Σ_*i*_*w*_*ij*_*o*_*i*_. The binary activation variable *o*_*j*_ represents unit activation and is related to the current through the WTA dynamics described in Eq. 2. This mechanism selects the unit receiving the maximum current at each hypercolumn and activates it. We introduce intrinsic adaptation as a mechanism controlled by the variable *a* in Eq. 3 to induce pattern deactivation. *dξ* represents additive white noise with variance *σ*. An extra current *I*_*j*_(*t*) is used to model external input into the system. For the sake of generality, it is important to stress that our current based population model is equivalent to a rate-based formalism as shown in Miller and Fumarola (2012).

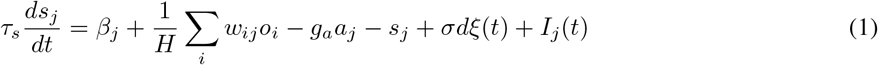

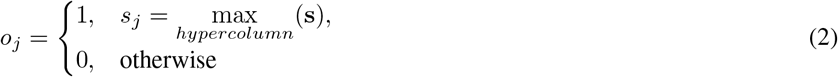

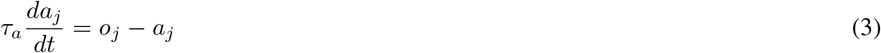

It has long been recognized that an attractor model with asymmetric connectivity produces sequential dynamics (Amit, 1992). In that vein, we explain now how an asymmetric connectivity matrix coupled with the dynamics of our model brings about sequential activity.

In Fig. 2A we show a case of successful sequential recall in the network with the connectivity matrix depicted in Fig. 2D. Here we handcrafted the connectivity matrix to illustrate the unfolding of the following dynamics. Once the first pattern gets activated (*o*_*i*_=1) as a result of an external cue (current input *I*(*t*) to all the units belonging to the pattern) the adaptation current *a*_*i*_ depicted in Fig. 2B starts growing and, in consequence, the self-excitatory current *s*_*i*_ becomes smaller. At some point, the self-excitatory current *s*_*i*_ is going to become weaker than the feed-forward current *s*_*i*+1_ which the next pattern in the sequence is receving. Then, the competitive WTA mechanism mediates the activation of the next pattern (*o*_*i*+1_ = 1) and suppresses the current one (*o*_*i*_) by competition. These dynamics are self-sustained and the cycle repeats until the end of the sequence. We depict the profile of such transitions in Fig.2C. The total time that the pattern stays activated is defined as the persistence time *T*_*per*_ (as used in van Hemmen et al. (1991)) and depends on the interplay between the connectivity matrix, the bias term and the adaptation.

**Table 1:** Relevant parameters and quantities.

| Symbol | Name | Values |
| --- | --- | --- |
| $\tau_s$ | Synaptic time constant | 10 *ms* |
| $\tau_a$ | Adaptation time constant | 250 *ms* |
| $g_a$ | Adaptation gain | 0 – 2.5 (units of $w$, control) |
| $\tau_{z_{pre}}$ | Pre synaptic z-filter time constant | 5 – 150 *ms* |
| $\tau_{z_{post}}$ | Post synaptic z-filter time constant | 5 *ms* |
| $\tau_p$ | Probability traces time constant | 5 *s* |
| $\sigma$ | Standard deviation of $s$ values | 0 – 3 |
| $T_{per}$ | Persistence time | 50 – 3000 *ms* (controlled) |
| $T_p$ | Pulse time | 100 *ms* |
| $\Delta T_p$ | Inter Pulse Interval (IPI) | 0 *ms* |

**Figure 2:**
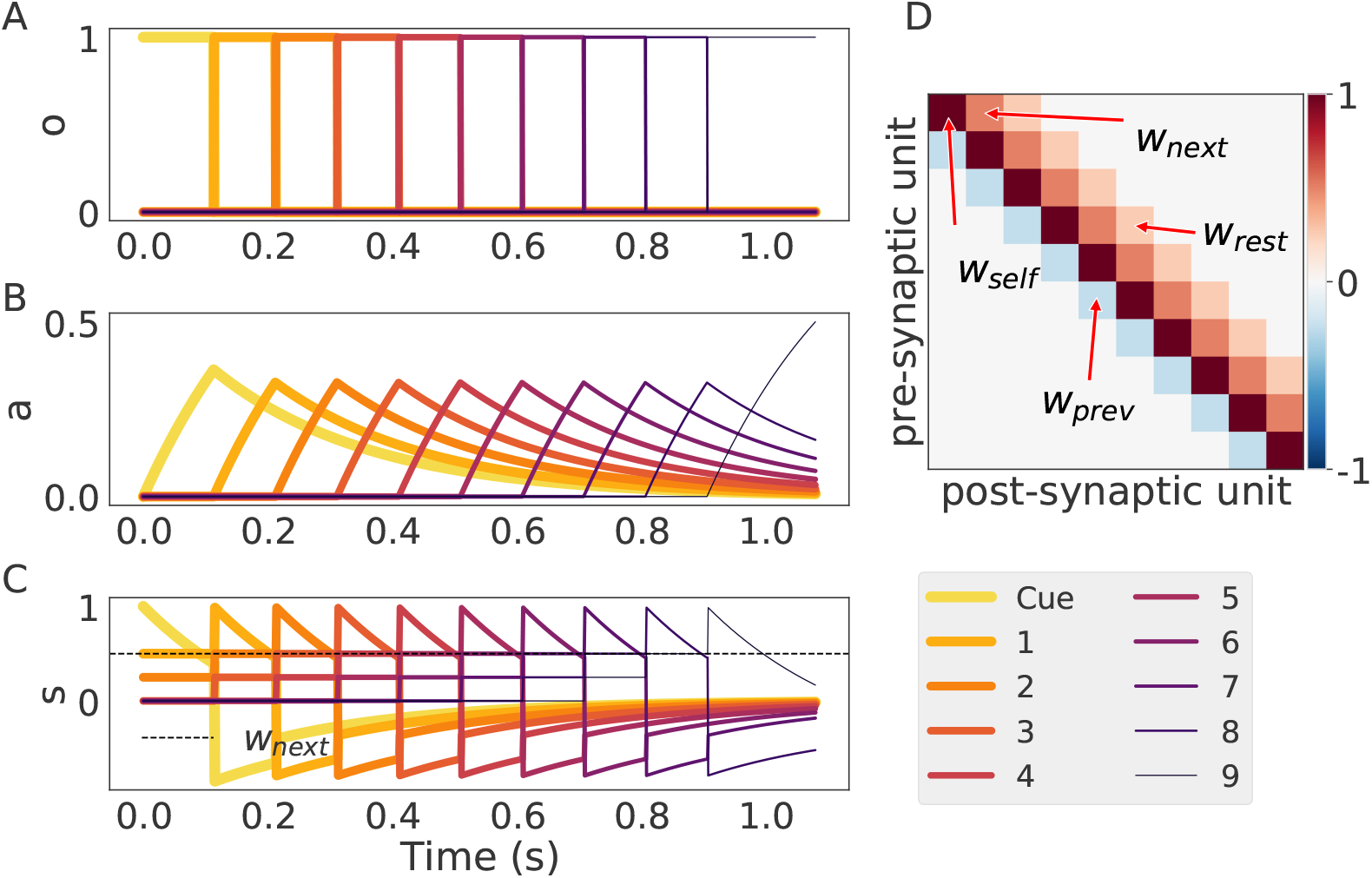
An instance of sequence recall in the model. (A) Sequential activity of units initiated by the cue. (B) The time course of the adaptation current for each unit. (C) The total current *s* (note that this quantity crossing the value of *w*_*next*_*o* depicted here with a dotted line) marks the transition point from one pattern to the next. (D) The connectivity matrix where we have included pointers to the most important quantities *w*_*self*_ for the self-excitatory weight, *w*_*next*_ for the inhibitory connection to the next element, *w*_*rest*_ for the largest connection in the column after *w*_*next*_ and *w*_*prev*_ for the connection to the last pattern that was active in the sequence.

### 2.2 Persistence time

Two important characteristics of sequence dynamics are the order in which the patterns are activated (the serial order) and the temporal structure of those activations (the temporal order) (Dominey and Ramus, 2000). In our model the serial order is determined by the differential connectivity between the current activated pattern and all other patterns. In general, the next pattern activated will be the one for which the quantity Δ*w*_*next*_ = *w*_*self*_ – *w*_*next*_ is smaller. The persistence time or temporal information of the sequence on the other hand is determined by the interplay between the connectivity of the network and the dynamical parameters of the network. We now proceed to characterize this relationship analytically. From the deterministic trajectories (see Appendix A) we can find the time point at which the currents from two subsequent units are equal: *s*_*i*_(*t*) = *s*_*i*+1_(*t*). Solving for *t* we determine the persistence time, *T*_*per*_ for each attractor determined with the expression in Eq. 4.

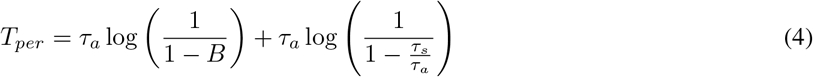

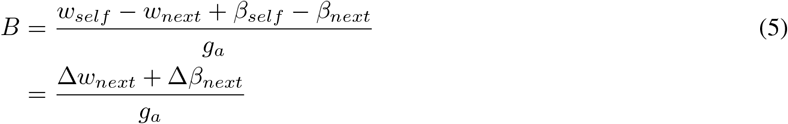

The parameter B in Eq. 5 condenses information regarding the connectivity *w*, bias terms *β*, and adaptation strength *g*_*a*_. From Eq. 4 we can infer that *T*_*per*_ is defined only for 0 < *B* < 1. This sets the conditions for how the weights, bias and external input interact with the adaptation parameters in order for the sequence to be learned and recalled. The straightforward interpretation for *B* < 1 is that the adaptation has to be strong enough to overcome the effects of the other currents, while *B* > 0 sets the connectivity conditions for sequence recall to occur (*w*_*self*_ > *w*_*next*_). As illustrated in Fig. 3A *T*_*per*_ is small for *B* ≈ 0 and diverges to infinity as *B* ≈ 1. This facilitates the interpretation of *B* as a unitless parameter whose natural interpretation is the inverse of transition speed, as shown in the examples provided in Fig. 3B-C.

**Figure 3:**
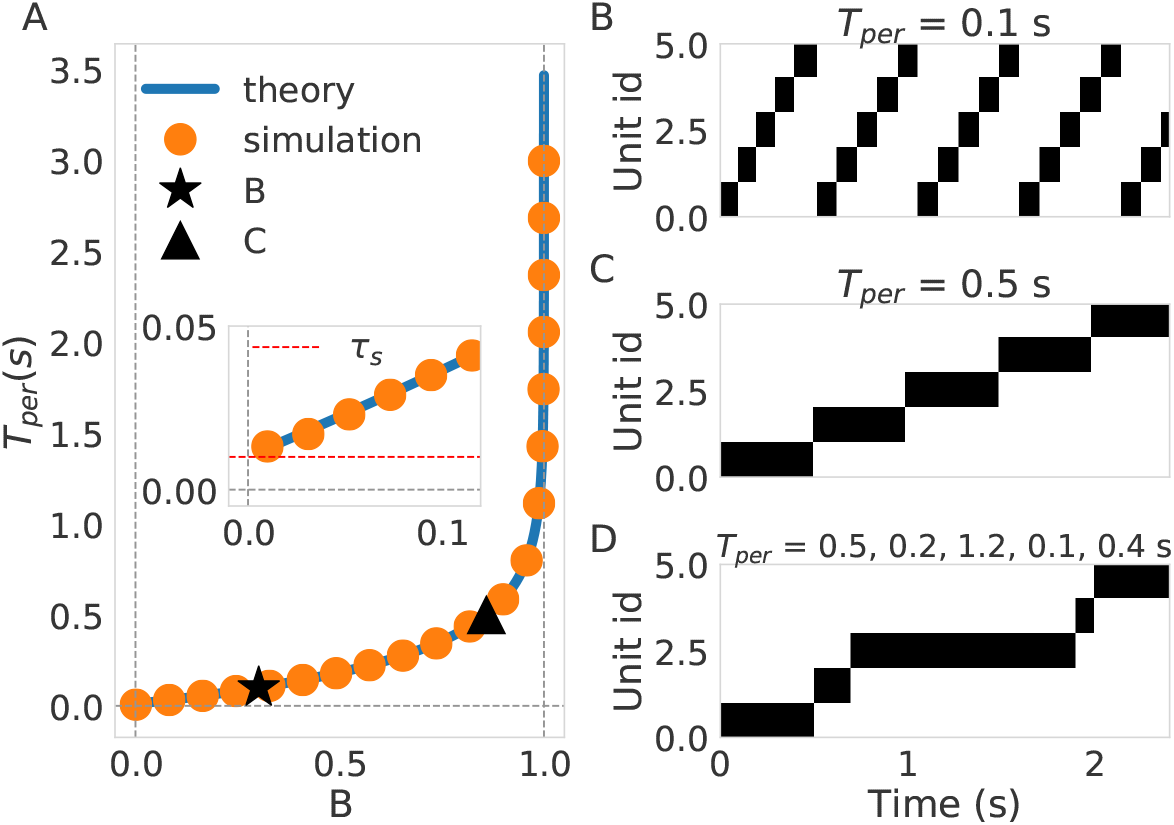
Systematic study of persistence time *T*_*per*_. (A) *T*_*per*_ dependence of B. The blue solid line represents the theoretical prediction described in Eq. 4 and the orange bullets are the result of simulations. Inset depicts what happens close to *B* = 0 where we can see that the lower limit is the time constant of the units *τ*_*s*_. (B) An example of sequence recall where *T*_*per*_ = 100 *ms*. This example corresponds to configuration marked the black star in (A). (C) example of sequence recall with *T*_*per*_ = 500 *ms*. This example corresponds to the configuration marked with a black triangle in (A). (D) Recall of a sequence with variable temporal structure (varying *T*_*per*_. The values of *T*_*per*_ are 500, 200, 1200, 100, and 400 ms respectively.

Controlling the individual persistence times of different patterns (the temporal structure) through short-term dynamics has been discussed previously in the literature (Veliz-Cuba et al., 2015). In our network the temporal structure of the sequence is also controlled by the adaptation dynamics. We illustrate this in Fig. 3D where by choosing specific values for the adaptation gain, *g*_*a*_, precise control of the *T*_*per*_ is achieved for every attractor.

For illustration purposes the formula in 4 is given for the case of orthogonal patterns and one hypercolumn. In the general case with more than one hypercolumn it is possible that not all transitions in a pattern (in different hypercolumns) occur at the same time. Moreover, as we recall sequences with non-repeating elements the adaptation effects are not specified. A full treatment that handles both the modular effects of non-overlapping elements and adaptation effects is given in Appendix A.

### 2.3 Learning

So far we have shown that our model can support sequence recall and control of the temporal structure through the adaptation dynamics. We now show that if the network is subject to the right spatio-temporal input structure then associative Hebbian learning is sufficient to induce the learning of the asymmetric connectivity structure characteristic of sequence recall (Amit, 1992). Based on previous work (Tully et al., 2016) we use the the BCPNN learning rule in its incremental on-line version (Sandberg et al., 2002) with learning mediated through asymmetric synaptic time traces. The version of the BCPNN learning rule presented is an adaptation of the discrete learning rule presented in (Lansner and Ekeberg, 1989) to a continuous setting.

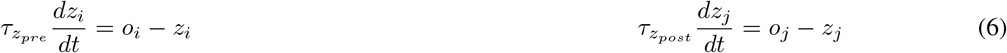

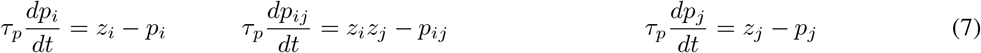

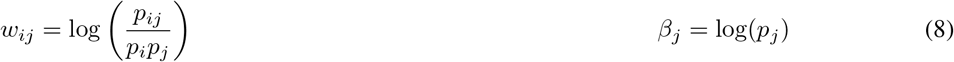

In the spirit of associative learning the BCPNN rule sets positive weights of recurrent connections between units that statistically tend to co-activate and creates inhibitory connections (negative weights) between those that do not. This is reflected in Eq. 8, where the connections are determined with a logarithmic ratio between the probability of co-activation (*p*_*ij*_) and the product of the activation probabilities (*p*_*i*_ and *p*_*j*_). Note that if the events are independent the weight between them is zero (*p*_*ij*_ = *p*_*i*_*p*_*j*_). Nevertheless, basic associative learning can only bind units that are active simultaneously. In order to bind units that are not simultaneously active in time we need an extra mechanism of temporal integration (Amit, 1992). To overcome this we combine the BCPNN learning rule with the introduction of the z-traces in order to create temporal associations between units that are contiguous in time (Tully et al., 2014). The z-traces, defined in Eq. 6, which can be thought of as synaptic traces, are a low-passed filtered version of the unit activations *o* and dynamically track the activation as shown in the top of Fig. 4B. To approximate the probabilities of activation (*p*_*i*_ and *p*_*j*_) and co-activation (*p*_*ij*_) the z-traces are accumulated over time in agreement with Eq. 7 which implements an on-line version of the exponentially weighted moving average (EWMA). As illustrated in Fig. 4A, asymmetry in the connectivity matrix arises from having two z-traces, a pre-synaptic trace with a slow time constant 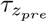 and a fast post-synaptic trace with a fast time constant 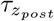 (Tully et al., 2016). In short, the z-traces work as a temporal proxy for unit activation that allow us to use the probabilistic framework of the BCPNN rule to learn the sequential structure of the input.

**Figure 4:**
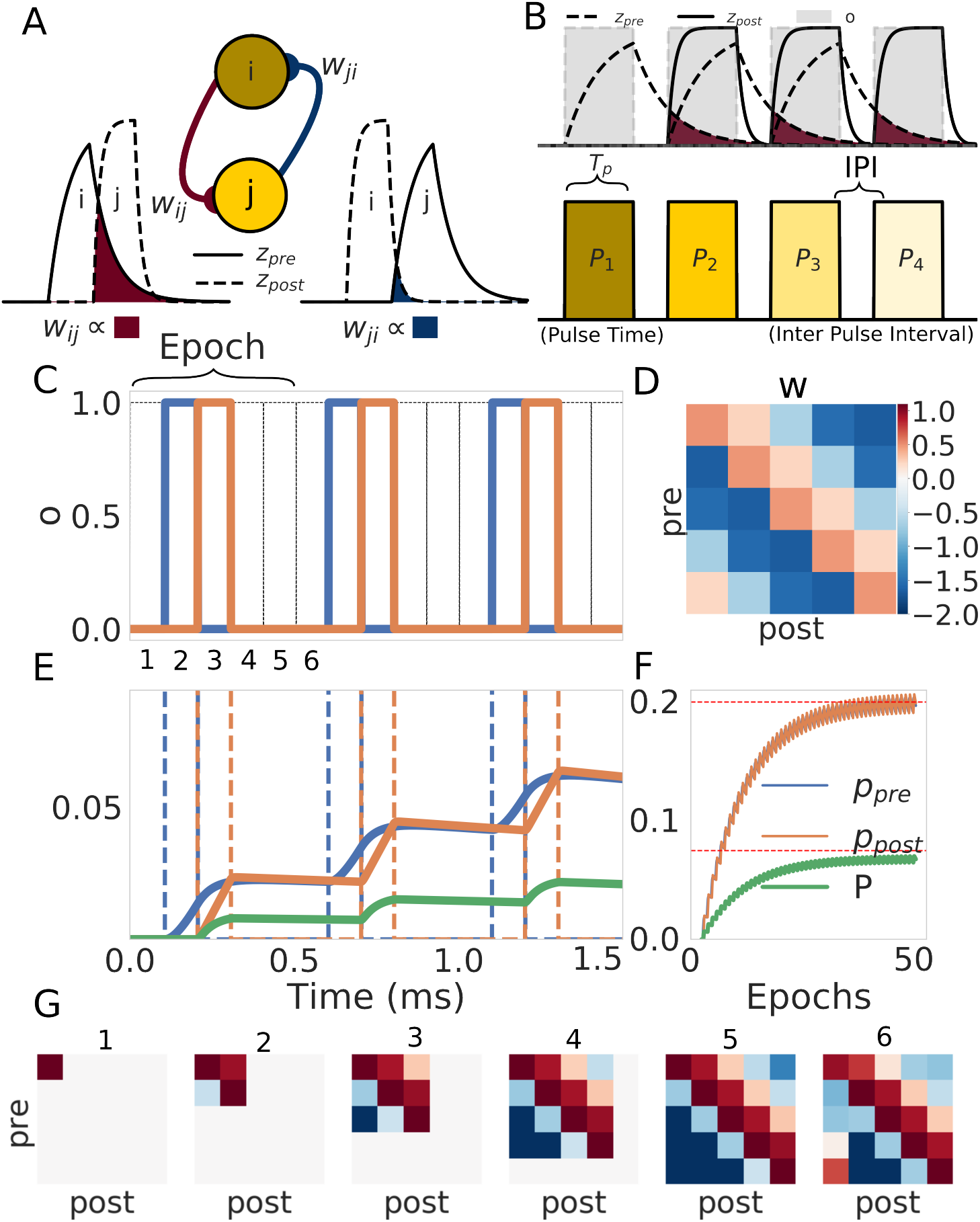
Sequence learning paradigm. (A) Relationship between the connectivity matrix *w* and the z-traces. The weight *w*^*ij*^ from unit *i* to unit *j* is determined by the probability of co-activation of those units which in turn is proportional to the overlap between the z-traces (show in dark red). The symmetric connection *w*_*ij*_ is calculated through the same process but with the traces flipped (here shown in dark blue). Note that the asymmetry of the weights is a direct consequence of the asymmetry of the z-traces. (B) Schematic of the training protocol. In the top we show how the activation of the patterns (in gray) induces the z-traces. In the bottom we show the structure of the training protocol where the pulse time *T*_*p*_ and the inter-pulse interval IPI are shown for further reference. (C) We trained a network with only five minicolums for illustration. The first three epochs (50 in total) of the training protocol are shown for reference. The values of the parameters during training were set to *T*_*p*_ = 100 *ms*, 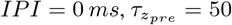 and 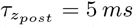. (D) The matrix at the end of the training (after 50 epochs). (E) Evolution of the probability values during the first three epochs of training. The probability values of the pre, post and joint probability evolve with every presentation. Note that the same color code is used in images C, E and F. (F) Long-term evolution of the probabilities with respect to the number of epochs. The values of the probability traces eventually reach a steady state. (G) Short-term evolution of the weight matrix at the points marked in the first epoch in C. Note that the colors are subjected to the same colorbar reference as in D.

The training protocol shown in Fig. 4B is driven by the temporal nature of the input and can be characterized by two quantities: the time that the network is exposed to a pattern (this is implemented by units being clamped through *I* in Eq. 1) called the pulse time, *T*_*p*_, and the time between the presentation of two patterns referred as the inter-pulse-interval (IPI). In the following we use a homogeneous training protocol where the values of the pulse time,*T*_*p*_, and the inter pulse interval, IPI, are the same for every pattern in the sequence.

The networks weights were learned using a training protocol where the patterns were presented sequentially for a number of epochs (50 epochs in the example illustrated in Fig. 4C-G). With every presentation of the stimuli the probability traces *p* absorb information (see Fig. 4E) slowly evolving to their steady state value (Fig. 4F). While the steady state weight matrix that results from training reveals asymmetric connectivity (Fig. 4D) the sequential structure of the input is learned as early as during the first epoch as can be observed in Fig. 4G. This demonstrates that the sequential structure of the input has been successfully learned by the BCPNN rule with the help of the z-traces.

We characterized the relationship between the connectivity matrix (*w*_*self*_, *w*_*next*_ and *w*_*prev*_) and the training protocol parameters (the pulse time *T*_*P*_, the inter-pulse-interval IPI and the two time constants of the synaptic traces 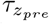 and 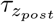). We summarize our findings and its relationship to the persistence time *T*_*per*_ in Fig 5. Longer pulse times *T*_*p*_ lead first to an increase in the value of *w*_*self*_ followed by its stabilization thereafter and to a decrease in the value of *w*_*next*_ (Fig. 5A). This can be explained by the fact that while the ratio between self co-activation and the total training time remains more or less constant (stabilizing *w*_*self*_) the co-activation between units becomes a smaller portion of the whole training protocol effectively reducing the estimating of *p*_*ij*_ (making *w*_*next*_ smaller). In consequence the rate of *T*_*per*_ growth becomes constant with longer pulse times *T*_*p*_ giving a logarithmic encoding of time (Fig. 5D). In contrast, longer inter-pulse-intervals lead to monotonic increments and decrements in *w*_*self*_ and *w*_*next*_ respectively (Fig. 5B). The reason for this is that a longer inter pulse intervals bring about an overall longer training protocol and after the co-activation of the units cease *p*_*i*_*p*_*i*_ decreases further than *p*_*ii*_ leading to a larger *w*_*self*_. *w*_*next*_, in the other hand, is rendered smaller by longer inter-pulse-intervals as a consequence of the unit’s activations begin further apart in time. It follows that *T*_*per*_ increases faster with larger IPIs as both *w*_*self*_ and *w*_*next*_ separate farther and farther with growing inter pulse intervals (Fig. 5E). The effect of the z-filters time constant *τ*_*z*_ in the weights can be described as diminishing the difference between *w*_*self*_ and *w*_*next*_ (5C). The results can be explained by interpreting the effect of increasing 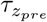 as spreading more and more the activation in time rendering the co-activations less meaningful overall (co-activation probability drops). This results in a diminishing value of *T*_*per*_ as the difference between weights Δ*w*_*next*_ drops with larger values of 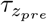 (Fig. 5F). Note here that the point at which 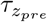 becomes larger than 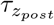 (marked with a dashed red line) coincides with *w*_*next*_ becoming larger than *w*_*back*_ as we should expect. The reasoning for *w*_*pre*_ is analogous to that of *w*_*next*_ with the only difference in synaptic time constant (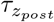 instead of 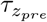).

**Figure 5:**
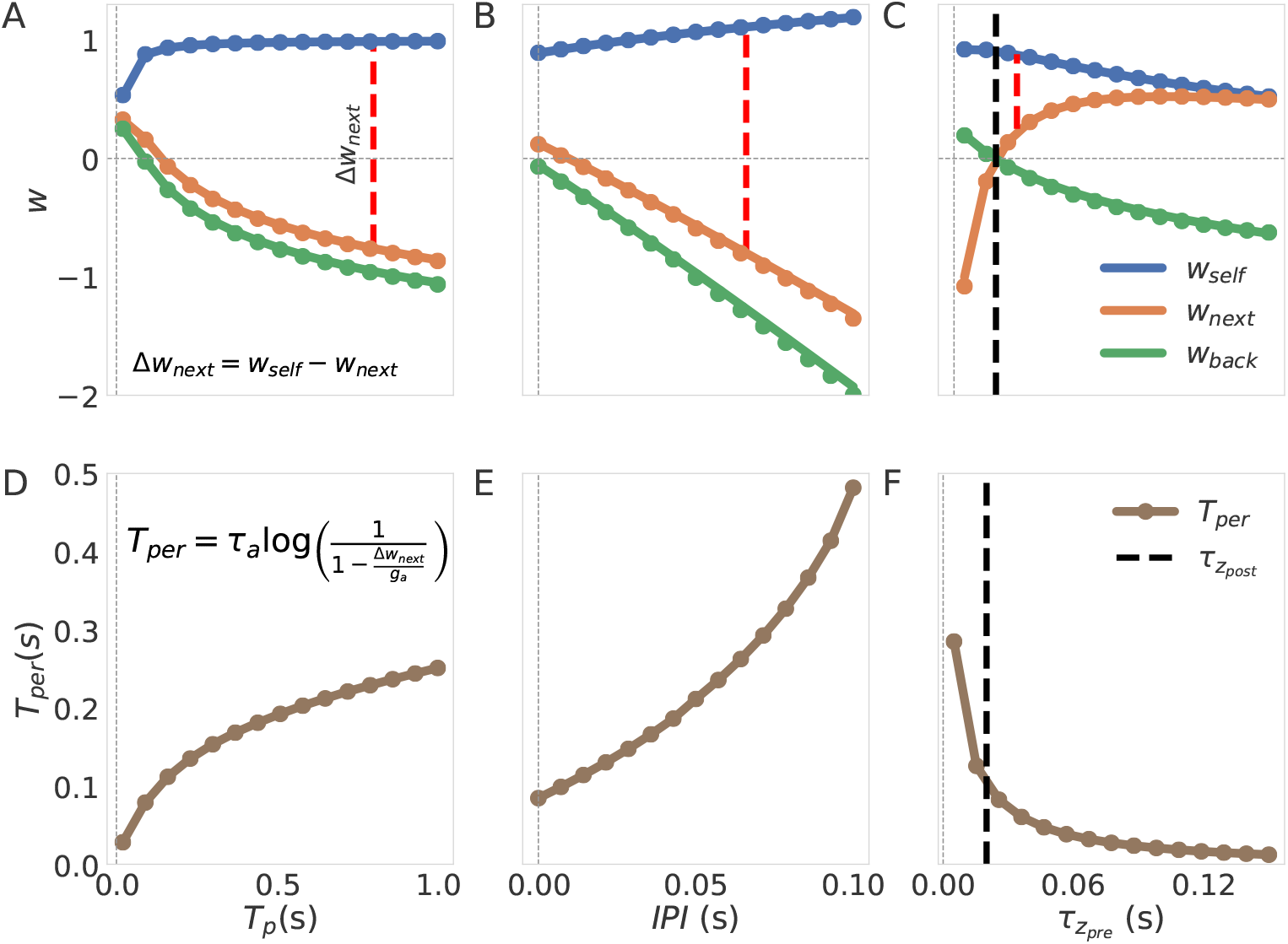
Characterization of the connectivity weights *w*_*self*_, *w*_*next*_ and *w*_*prev*_. We also show the effects of training in the persistence time *T*_*per*_ of the attractors. The equation on the inset in D relates *T*_*per*_ to Δ*w*_*next*_ = *w*_*self*_ *w*_*next*_ which we show as dashed red lines in each of the top figures (note that here Δ*β* = 0 as we trained with an homogeneous protocol). When the parameters themselves are not subjected to variation their values are: 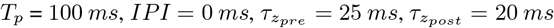 for all the units. (A-C) Show how the weights depend on the training parameters *T*_*p*_, inter pulse interval and 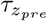, respectively, whereas (D-E) illustrate the same effects on *T*_*per*_. Here we are providing the steady state values of *w* obtained after 100 epochs of training.

We have shown so far that the temporal structure of the input determines the temporal structure of the recall (Fig5D-F). We now show that the inter-pulse-interval, IPI, can change the recall phase from a sequence regime where the patterns are tied in time (Fig. 6A) to a free attractor regime, where the patterns are learned independently (Fig. 6B). In general, to bridge a longer inter-pulse-interval, a longer 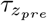 is required as illustrated in Fig 6C. The idea is that 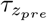 provides a temporal window of integration that links the patterns in time and the larger the window is, the longer are the inter-pulse-intervals that it can bridge.

**Figure 6:**
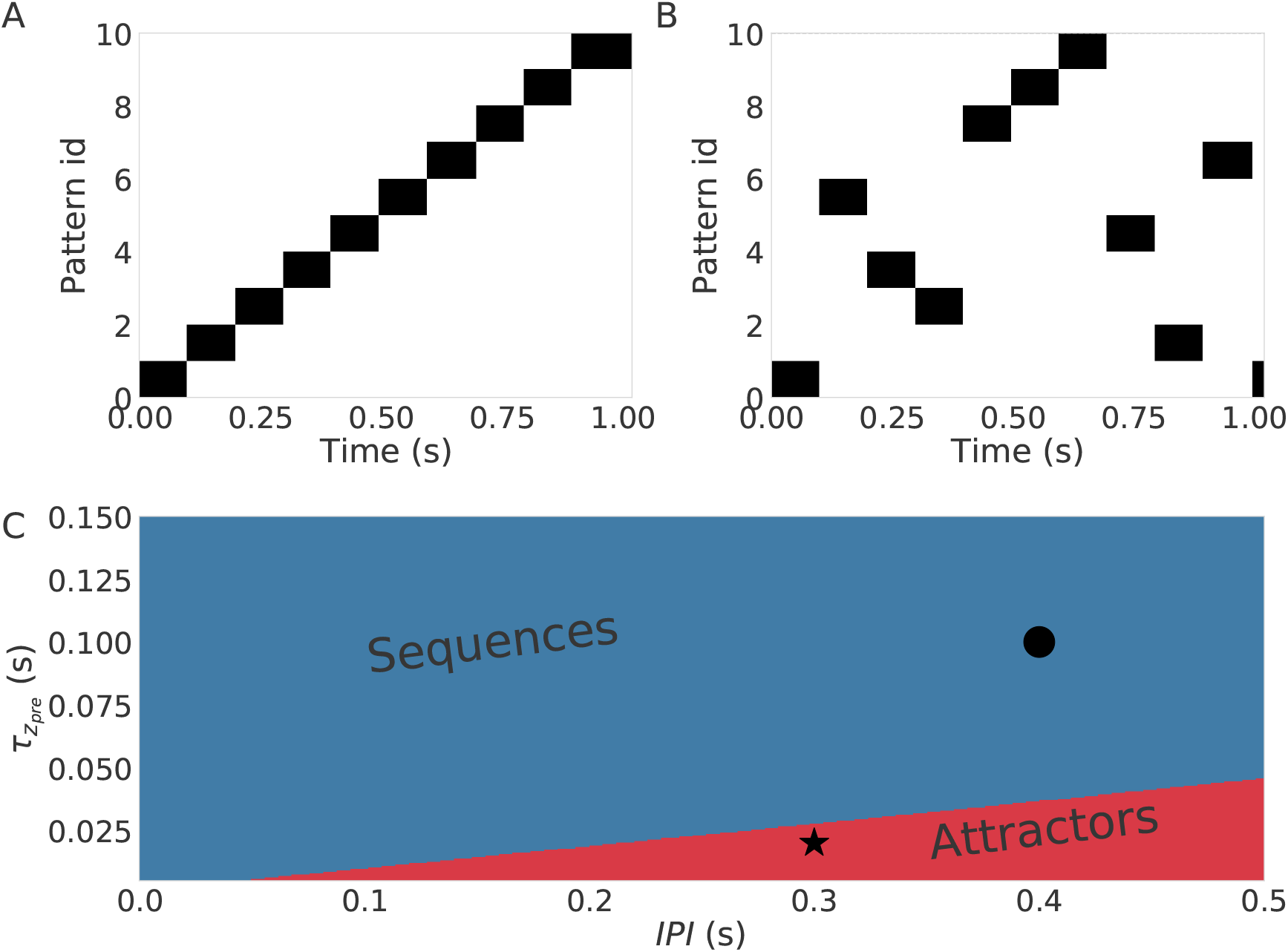
Transition from the sequence regime to the free attractor regime. (A) An example of a sequential (ordered) activation of patterns. (B) An example of an unordered chain of activations of patterns in the free attractor regime. (C) The two regimes (sequential in blue and free attractors in red) in the relevant parameter space spammed by 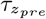 and inter pulse interval. The examples in (A) and (B) correspond to the black dot and the star, respectively.

### 2.4 Noise

We also tested whether sequence recall in the network was robust to noise by controlling the level of noise with the parameter *σ* in Eq. 1. Additive noise manifest itself in stochastic trajectories where pattern to pattern transitions happens earlier (Fig. 7A). This phenomenon is illustrated clearly with the red and purple lines in Fig 7A where compared to their deterministic counterparts (solid lines) the noisy trajectories (thin lines) make the transition as soon as the variations in *s* drive them under the transition point (*w*_*next*_ *o*). Therefore, the persistence time in a network operating in a noisy regime will be a stochastic variable (denoted *T*_*per,σ*_) whose mean will be lower than the persistence time *T*_*per*_ present in the deterministic regime. The mean value of *T*_*per,σ*_ decays systematically with increasing sigma and quickly converges to a common value independent of the value of *T*_*per*_ for the deterministic regime set by controlling *g*_*a*_ (Fig. 7B). To examine whether a sequence with lower values of *T*_*per*_ is less likely to be recalled correctly under the influence of noise we cued the sequence 1000 times for every value of *σ* and constructed the success rate vs noise profile shown in Fig. 7C where we observe that the success rate is identical for different values of *T*_*per*_. We conclude that *T*_*per*_ has no effect in how sensitive the recall process is to noise thus facilitating the study of the effect of noise in the system by enabling us to control *T*_*per*_.

**Figure 7:**
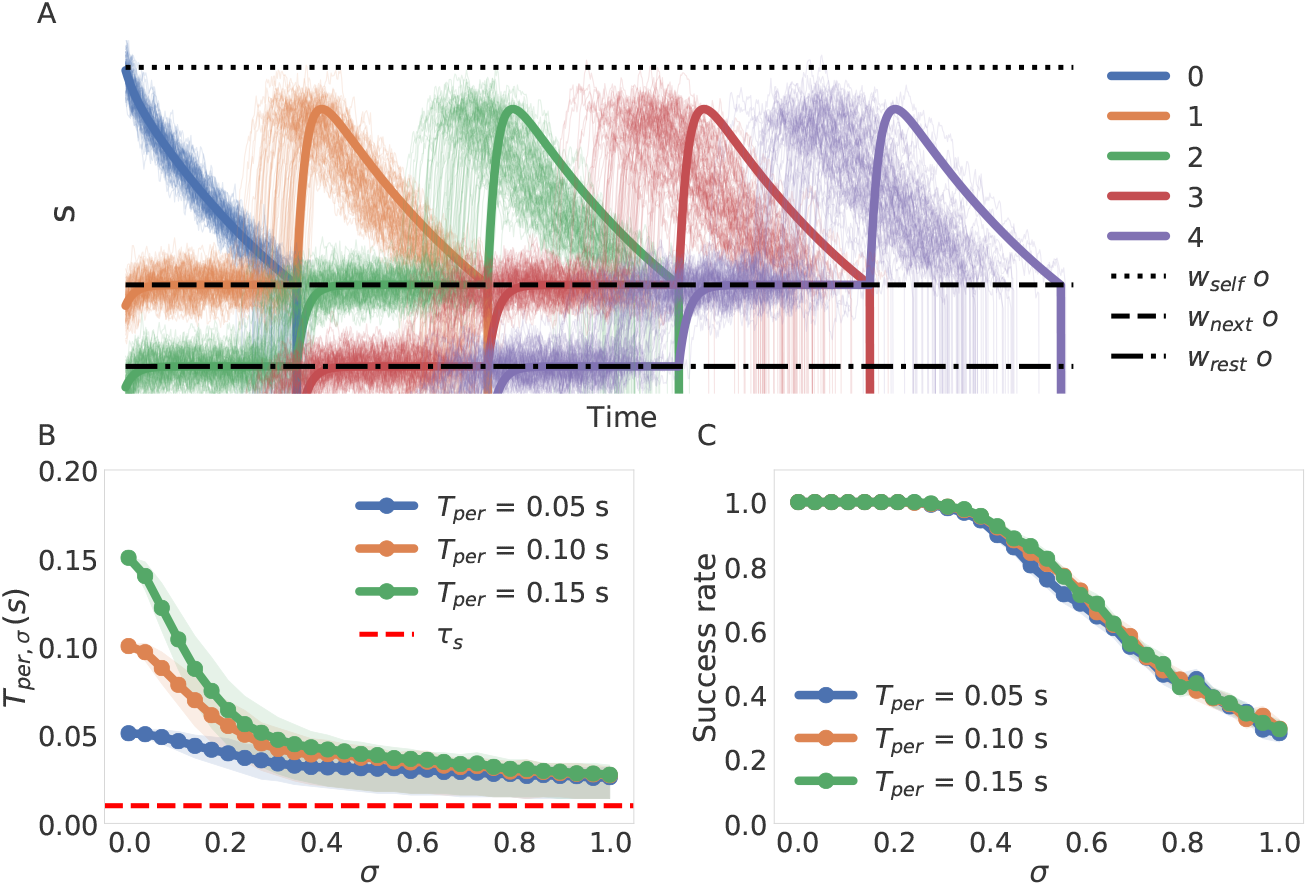
Effects of noise reflected in current trajectories and persistence times. (A) An example of current trajectories subjected to noise. The solid lines indicate the deterministic trajectories the system would follow in the zero noise case. In dotted, jagged and dashed lines we depict the currents induce *w*_*self*_, *w*_*next*_ and *w*_*rest*_ for reference. (B) Change in the average of the actual value of *T*_*per*_ for different levels on noise. We Shaded the area between the 25th and the 75th percentile to convey and idea of the distribution for every value of *σ* (C) Success rate vs noise profile dependence on *T*_*per*_. We ran 1000 simulations of recall and present the ratio of successful recalls as a function of *σ*. Confidence intervals from the binomial distribution are too small to be seen.

Next we systematically characterized the sensitivity of the network to noise as a function of the training parameters by calculating *σ*_50_ (see Methods). We illustrate the nature of *σ*_50_ in Fig. 8A, please note that a larger *σ*_50_ implies a system which is less sensitive to noise and vice versa. Calculating *σ*_50_ for different values of *T*_*p*_ we conclude that the network becomes less sensitive to noise with longer values of *T*_*p*_ as shown in 8B. This can be explained by the fact that training with longer pulses increases the distances between the weights (and therefore the distance between the currents) as previously shown in Fig. 5A. We can see the same effect by increasing the inter pulse interval in Fig. 8C where the separation of weights produced by longer inter pulse intervals leads to a similar outcome. The opposite effect is observed with longer values of 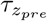 where the system becomes more sensitive with longer values of 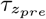 as shown in 8D. We can appeal again to the structure of the weights in Fig. 5C to explain these results as an outcome of the weights and therefore the current being less differentiated among themselves leading to failures in sequence recall.

**Figure 8:**
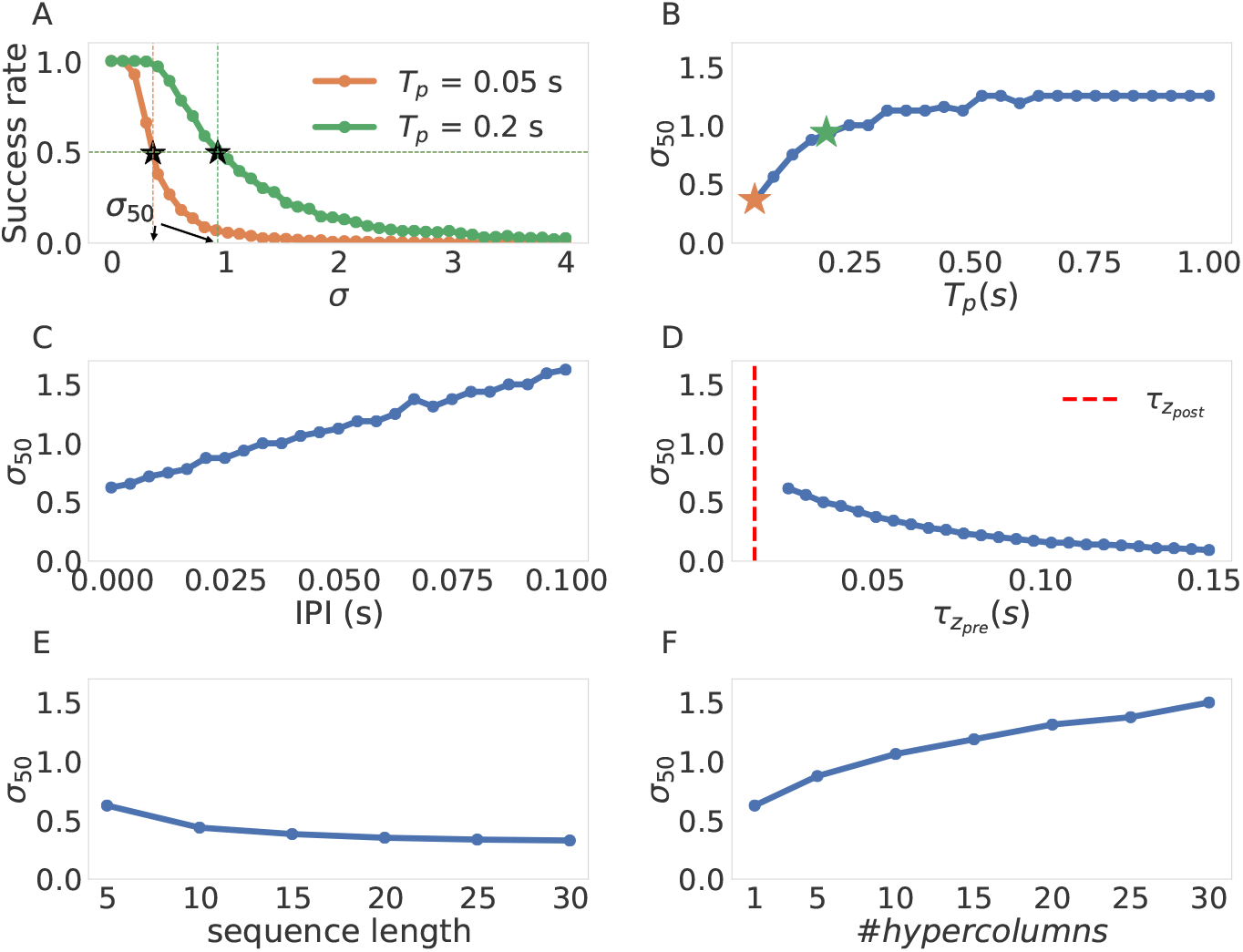
Sensitivity of network performance to noise for different parameters. The base reference values of the parameters of interest are: 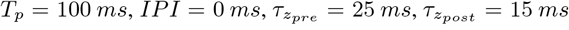, sequence length = 5, #hypercolumns = 1. (A) Two examples of the success vs noise profiles (*T*_*p*_ = 50 *ms*, 200 *ms*). The value of *σ*_50_is indicated in the abscissa for clarity, note that smaller *σ*_50_implies a network that is more sensitive to noise (the success rate decays faster). (B) *σ*_50_ variation with respect to *T*_*P*_. We also indicate the *σ*_50_ for the values of *T*_*p*_ used in (A) with stars of corresponding colors.(C) *σ*_50_variation with respect to the inter pulse intervals. (D) *σ*_50_ variation with respect to the value of 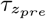. (E) *σ*_50_ variation with respect to sequence length. (F) *σ*_50_ variation with respect to the number of hypercolumns.

We also report two relevant noise effects not related to the connectivity. First, we show in Fig. 8E that the network becomes more sensitive to noise for longer sequences. This can be explained by considering each pattern-to-pattern transition as a possible point of failure. Naturally, adding more links to the chain makes the recall of the sequence more likely to fail at some point (i.e. not recall all patterns in the right order). Finally, in Fig 8F we observe a scaling effect in how robust the network is with the number of hypercolumns. This can be explained using the fact a network with more hypercolumns posses a higher degree of recurrent connectivity. Every time there is a mis-transition in any of the units the recurrent connectivity channels the currents of the units where the transition occurred correctly as an error correction mechanism assuring the successful completion of the sequence more often than not. In a more abstract language the more hypercolumns the network possess, the less likely it is for enough transitions to occur such that the network state is pushed out of the basin of attraction of the next pattern. Therefore, the more hypercolumns the network possess, the more robust it is to noise and hence the observed scaling.

### 2.5 Overlapping representations and sequences

Previous work with attractor models has shown that it is possible to store attractor states with overlapping representations (i.e. patterns that shared a unit activation in some hypercolumns) (Meli and Lansner, 2013; Sandberg et al., 2002). We test here whether our network is able to store and recall overlapping patterns successfully when they belong to sequences and are recalled as such. This is desirable to increase the storage capacity of our network and to enrich the combinatorial representations that our network can process.

Our aim is to characterize the capabilities of our network to store and successfully recall sequences containing patterns with some degree of overlap. As sequences can contain more than a pair of overlapped patterns we propose the following two parameters as a framework to systematically parameterize the problem: 1) the first parameter quantifies the level of overlap between the representation of two patterns and is therefore a spatial measure of overlap, we call this parameter representation overlap. 2) the second parameter is a temporal metric of overlap and quantifies how many patterns between two sequences possess some degree of representational overlap; we call this parameter sequential overlap. A schematic illustration of the general idea is presented in Fig. 9A1, where the two parameters, the representational overlap and the sequential overlap, are shown in black and grey, respectively. To be more precise, the representational overlap between two patterns is defined as the proportion / ratio of hypercolumns that share units between the two patterns. We define the sequential overlap between two sequences as the number of patterns in the sequences that possess some degree of overlap (e.g. in Fig. 9A1 the sequential overlap is 4). In order to illustrate these concepts we present a detailed example in Fig. 9B. The example consists of two six-pattern sequences (i.e. of length six) whose patterns are distributed over three hypercolumns (for example, the first pattern *P*_1*a*_ of sequence a consists in the activation of the unit 10 in each of the three hypercolumns). The two sequences have two pairs of patterns that have some degree of overlap (pairs *P*_3*a*_ − *P*_3*b*_ and *P*_4*a*_ − *P*_4*b*_) and therefore the two sequences have a sequential overlap of 2 as indicated by the gray area in Fig 9B. If we look at patterns *P*_3*a*_ = (12, 3, 3) and *P*_3*b*_ = (3, 3, 3) we can observe that they have the same unit activation in the last two hypercolumns (hypercolumns 2 and 3) and therefore the pair has a representational overlap of 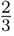. The units in the hypercolumns responsible for the representational overlap between the pair are highlighted in black in Fig. 9B. Note that the representational overlap is a parameter between 0 and 1, whereas the sequential overlap is an unbounded parameter as sequences can be arbitrarily long.

**Figure 9:**
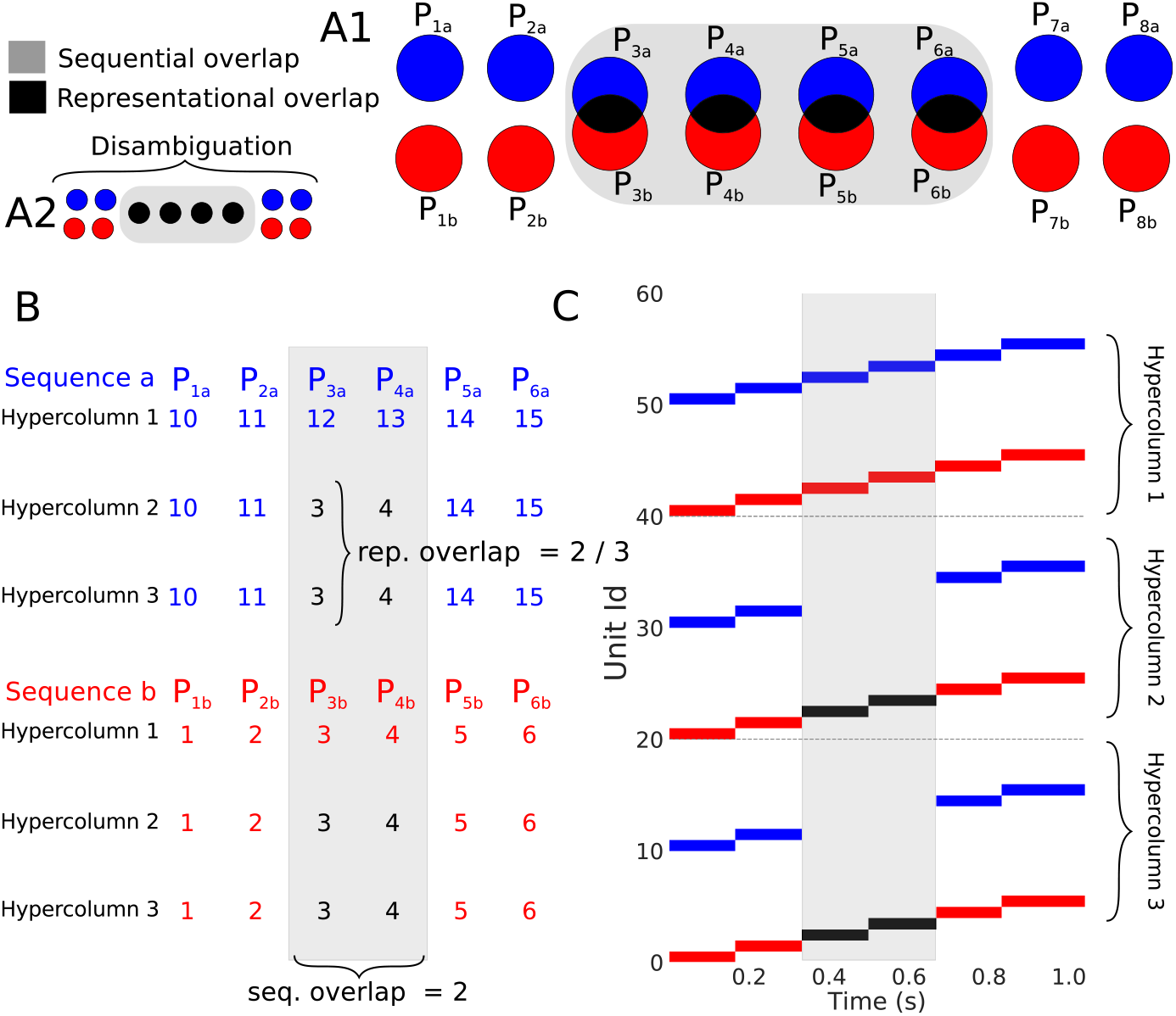
Overlapping representations and sequences. (A1) Schematic of the parameterization framework. Black and gray stand for the representational overlap and the sequential overlap respectively (see text for details) (A2) Schematic of the sequence disambiguation problem. (B) An example of two sequences with overlap. Here each row is a hypercolumn and each column a pattern (patterns *P*_1*x*_, *P*_2*x*_, *P*_3*x*_, *P*_4*x*_, *P*_5*x*_, and *P*_6*x*_). The single entries represent the particular unit that was activated for that hypercolumn and pattern. (C) The superposition of the recall phase for the sequences in (B). Each sequence recall is highlighted by its corresponding color. We can appreciate inside the gray area that the second and third hypercolumns (sequential overlap of 2) have the same units activated (depicted in black). This reflects the fact those patterns have a representational overlap of 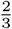 (two out of three hypercolumns).

The limit case when representational overlap is equal to 1 is the domain of sequence disambiguation. We show a schematic of the disambiguation problem in Fig. 9A2 where a representational overlap of 1 can be interpreted as the equivalence of both patterns in the sequential overlap section. In this regime the sequential overlap corresponds to the size of the disambiguation window that the network has to bridge to correctly disambiguate the sequence (i.e. ending in *P*_8_*a* if you started in *P*_1_*a* in Fig 9A2). Solving sequence disambiguation in the most strict sense requires the network to be able to store the contextual information required to solve correctly the bifurcation at the end of the overlapping section. That is, the network requires to hold the information of what pattern was activated before the disambiguation window for as long as the time it takes for the sequential dynamics to reach forking point.

In general we should expect that sequences with higher representational and sequential overlaps would be harder to process for the network. To characterize these difficulties systematically we tested for correct sequence recall for sequences in the zero noise condition for all the possible combinations of representation overlap as well as sequential overlap that the network allowed. As can be see in Fig. 10A the network can successfully recall overlapping sequences over a wide range of sequential and representational overlaps. The exception to this is the disambiguation regime in top of Fig 10A where we see a failure to recall both sequences when overlapped patterns are identical. Next we studied the recall of sequences with overlapping patterns in the presence of noise. First, we examined the dependence of the success rate on the noise level for a wide array of sequential and representational overlaps (1, 2, 3 and 4 in Fig. 10A). The results, as shown by the curves in Fig 10B, illustrate that the success rate vs noise profiles are very similar despite different degrees of sequential and representational overlap. Second, for a fix value of representational overlap (0.5), we calculated *σ*_50_ for all the possible values of sequential overlap (green horizontal line in Fig. 10A). We also calculated the values of *σ*_50_ for a fix value of sequential overlap (5) and all the possible values of representational overlap (blue vertical line in Fig. 10A). The results (Fig. 10C,D) show that the network is robust to noise across the spectrum of possible overlaps except when we get close to the sequence disambiguation regime (right part of Fig 10D), where the network becomes more sensitive. Those results together suggests that our neural network can consistently recall sequences correctly over a broad set of overlap conditions.

**Figure 10:**
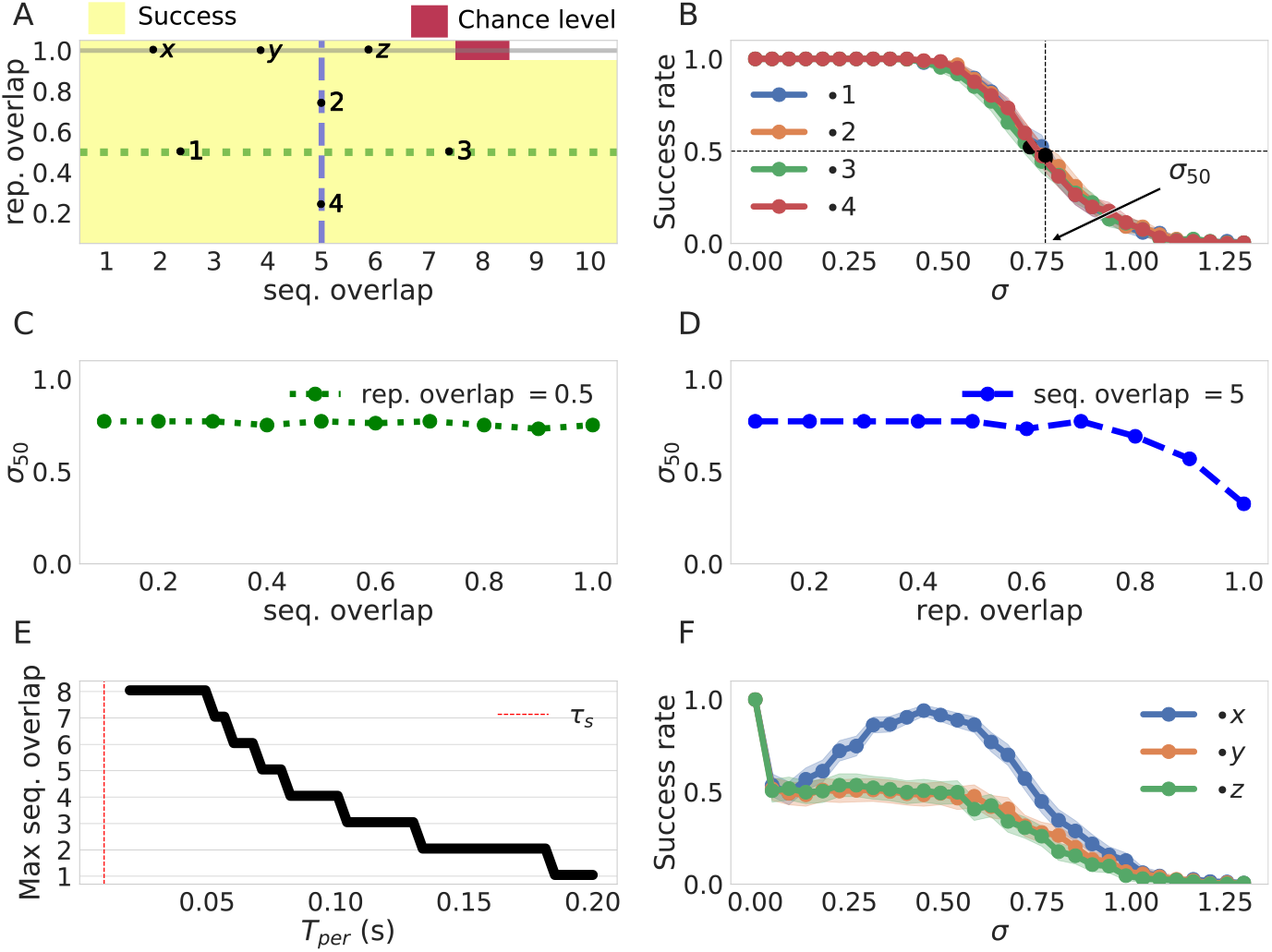
Sequence recall performance for different overlap conditions. The base line values of the parameters of interest are 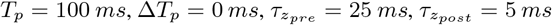 sequence length = 10, # hypercolumns= 10 and *T*_*per*_ = 50 *ms*. (A) Success rate for pairs of two sequences with different sequential and representation overlaps. We show here the performance over the parameter space. Success here is determined by correct recall of both sequences. (B) Success rate vs noise level for the sequences with configurations marked as 1, 2, 3, 4 in A. The values of *σ*_50_ are marked as an illustrations for the calculations below. (C) *σ*_50_ as a function of the sequential overlap. The values of *σ*_50_ are calculated over the sequences with configurations given in the green horizontal line in A. (D) *σ*_50_ as a function of the representation overlap. The values of *σ*_50_ are calculated over the sequences with configurations given in the blue vertical line in A. (E) max disambiguation as a function of *T*_*per*_. The network loses disambiguation power with long lasting attractors as the memory of the earlier pattern activation reflected in the currents fades. (F) Success rate vs noise profile in the disambiguation regime. The three curves correspond to overlapping sequence configurations marked with x,y, and z in A. Shaded areas correspond to 95% confidence intervals.

In the disambiguation regime with no noise (gray line in Fig. 10A) the network is able to solve the disambiguation problem successfully up to disambiguation windows of size 8. The disambiguation capabilities of the network are due to memory effects on the dynamics (here capacitance effects mediated by *τ*_*s*_). In fact, we show in Fig. 10E that the longer the persistence times (and therefore the more time for the memory to fade) the smaller is the disambiguation window that the system can resolve. Contrary to the results above the network is brittle in the sequence disambiguation regime. In particular, the success rate decays extremely fast in the presence of noise as show in Fig 10F. However, an interesting resonance phenomena occurs for low sequential overlaps (blue curve) where the success rate actually increases with noise. This can be explained with the fact that the noise effectively reduces the mean persistence time *T*_*per,σ*_ (as shown before in Fig. 7B) which leads to the increased disambiguation power (c.f. 9E). to memory effects on the dynamics (here capacitance effects mediated by *t*_*s*_). In fact, we show in Fig. 10E that the longer the persistence times (and therefore the more time for the memory to fade) the smaller is the disambiguation window that the system can resolve. Contrary to the results above the network is brittle in the sequence disambiguation regime. In particular, the success rate decays extremely fast in the presence of noise as show in Fig 10F. However, an interesting resonance phenomena occurs for low sequential overlaps (blue curve) where the success rate actually increases with noise. This can be explained with the fact that the noise effectively reduces the mean persistence time *T*_*per,σ*_ (as shown before in Fig. 7B) which leads to the increased disambiguation power (c.f. 9E).

## 3 Discussion

We have evaluated a Hebbian-like BCPNN learning rule with asymmetrical temporal synaptic traces as a sufficient principle underlying robust sequence learning in an attractor neural network model. The results have revealed the potential of the network to successfully encode and reliably recall multiple overlapping sequential representations even in the presence of noise. In this context, we have systematically studied the effect of network modularity as well as the role of key temporal parameters of the synaptic learning rule. We have also stressed that our network has the capability to control the temporal structure of the sequential pattern recall by means of an intrinsic adaptation mechanism.

### 3.1 Previous work and biological context

Here we have followed the modelling philosophy aimed at distilling the architecture of the network to its essential characteristics that support and control the phenomenon of interest (sequence learning). In the previous models of particular relevance to our work, complex spike based dynamics and rich biological detail were promoted to provide insights into the biophysical underpinnings of sequence learning in the cortex (Tully et al., 2016) and as a model of memory consolidation (Fiebig and Lansner, 2017). While the aforementioned contributions provide a more direct mapping between biology and the network, our approach, which reduces the network to its essential characteristics, necessarily dilutes that mapping. Nevertheless, some key design principles emerging from biology are preserved. Below we discuss in more detail the main aspects of the relationship between the dynamical as well as structural properties in our network and the biological substrate that inspired them in the first place.

A general characteristic of cortical circuits is competition (Douglas and Martin, 2004). Competition is modelled in our network locally with a WTA mechanism but our results do not change qualitatively if a weaker soft-max mechanism is implemented instead (data not shown). Besides, Douglas and Martin (2004) suggested that such a competition mechanism could be implemented by basket or chandelier cells. In Tully et al. (2016) this computational principle was implemented by means of fast inhibitory basket cells with fixed connectivity and produced the same outcome. It is important to point out that the idea of using diverse forms of local competition to achieve pattern selection in sequence recall has been examined previously and extensively in the sequence learning literature (Mostafa and Indiveri, 2014; Murray et al., 2017; Byrnes et al., 2011).

Asymmetrical temporal traces have been proven successful to achieve the effect of sequence learning (Herz et al., 1989; Coolen and Gielen, 1988; Abbott and Blum, 1996; Lawrence et al., 2006; Veliz-Cuba et al., 2015; Pereira and Brunel, 2018). In our model we have utilized the temporal asymmetric z-traces as the basis of probabilistic learning with the BCPNN learning rule. The degree of asymmetry of the z-traces and its effects on the connectivity matrix have been studied through variations in 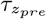 (Fig. 5C). In this framework lower values of 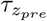 would correspond to fast AMPA dynamics (Holthoff et al., 2010) while longer values of 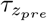 would correspond in turn to slower NMDA dynamics (Paoletti et al., 2013). Consistently with these observations, throughout this work we have restricted the values of 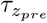 to the 5 – 150 *ms* range. A biological account of the z-traces and their connection to the biochemical cascades that underlie synaptic learning have been presented in a more detailed way by Tully et al. (2014).

It is important to point out that synaptic connections learned in our network with the BCPNN learning rule violate Dale’s law, i.e. projections emanating from the same unit can mediate both excitatory and inhibitory effects on the target units. To address this issue, we propose a different interpretation for positive and negative synaptic weights. In the former, they can be straightforwardly interpreted as the conductance between two units, whereas in the latter case we interpret them as a disynaptic connection through an inhibitory interneuron. The argument for the biological plausibility of this arrangement using double bouquet cells as the inhibitory interneurons in this architecture is developed furhter by Chrysanthidis et al. (2018).

### 3.2 Control of the temporal structure of the sequence

We have shown that the persistence time, *T*_*per*_, of our attractors can be quite effectively controlled through the use of the adaptation gain *g*_*a*_ and less effectively by means of the adaptation time constant *τ*_*a*_ (see Fig. 3 and Eq. 4). The range of *T*_*per*_ values for the attractor patterns in our network model is within the 10 *ms* and 3.5 *s* range. This in turn means that the duration of our sequences corresponds to the milliseconds to minutes interval (considering sequential lengths of 10 to 100). This range of values is consistent with the variation in sequence duration that Bhalla (2017) found for biological sequences in the hippocampus. While the mechanisms for temporal phenomena under the millisecond scale (inter-aural-scale, Carr and Konishi (1990)) and over the minute scale (circadian rhythms, Golombek et al. (2014)) are already well understood, the nature and origin of temporal phenomena at the intermediate time scales is still a matter of debate (Paton and Buonomano, 2018). We believe our work contributes to this debate by offering an intrinsic model of time (Ivry and Schlerf, 2008) capable of both, using the taxonomy of Paton and Buonomano (2018), the production and reproduction of temporal patterns within the discussed range.

In the work of Murray et al. (2017) the control of the temporal structure (control of *T*_*per*_) is accomplished by means of input from an external network. Although the ability of our network to control the temporal structure rests on internal mechanisms, we could also exploit external input for this purpose. By adding external input to our differential equation during recall and solving the resulting expression (see Appendix A) we obtain an expression for our parameter *B* in the following form *B* = (Δ*w*_*next*_ + Δ*β*_*next*_ + Δ*I*(*t*))(*g*_*a*_)^-1^ where Δ*I*(*t*) = *I*_*self*_ (*t*) – *I*_*next*_(*t*) is the differential input between the consecutive units in the sequence. By controlling this differential input, the persistence time of attractor states in a given sequence can be modulated. This could be used to build a framework where a generalist network learns the sequential structure of the input and a specialized control network adjusts the temporal structure of the sequence recall suitable for the task at hand.

### 3.3 Sequence Disambiguation and overlapping representations

Sequence disambiguation or using past context to determine the trajectory of a sequence has been deemed one of the most important problems that a sequence prediction network should solve (Levy, 1996). While some networks (Sussillo and Abbott, 2009; Rajan et al., 2016; Wang et al., 2017) have addressed the problem in their generality, their reliance on supervised learning and lack of biological plausibility remain a matter of concern. There have been a few attempts at the problem of sequence disambiguation in the attractor network framework but most of them rely on non-local learning rules or require an infeasibly large number of parameters (Fukushima, 1973; Guyon et al., 1988; Amit, 1992). Minai et al. (1994) proposed an alternative approach using the activity in a random network (what now is called a reservoir) as a source of context information for disambiguation. In their network, activity in the reservoir evolved in a path-dependent way, and inter-network connectivity between the disambiguation network and the reservoir conveyed the necessary information from the latter to the former thus allowing for successful disambiguation. While effective, such networks require another complete layer to keep a dynamical memory, an approach judged to be inefficient. To address this issue, context codes with less overhead have been proposed where, instead of a network, the state of a unit or a collection of units is determined by the dynamical history of the system and that state is then used for disambiguation (Sohal and Hasselmo, 1998; Samura et al., 2008). In our network, disambiguation can be achieved by building cell assemblies containing a subset of units that are preferentially connected to the subsequent assembly in the sequence. By preferential connectivity we mean that those units posses strong excitatory connections to the units of the subsequent pattern and strong inhibitory connections to the rest. To put it more concretely, the BCPNN learning rule, following its probabilistic nature, will ensure that the non-overlapping parts in a sequence are connected in such fashion by creating excitatory units between the units in the non-overlapping parts and the subsequent units in the sequence (as they are the only ones that actually appeared together) and strong inhibitory connections between the non-overlapping units and all the units belonging to any other pattern (as they never appeared together). In virtue of the aforementioned connectivity, activation of the units in the non-overlapping part of the assembly (context units) guarantees a transition to the subsequent (correct) pattern. As shown in Fig. 10D, the proposed mechanism is very robust to the size of the cell assembly that gets connected preferentially (the non-overlapping part); degradation of the performance under noise only becomes evident when the size of the context code becomes less than 20% of the cell assembly. This is consistent with some experimental evidence of neurons in the hippocampus that fire in such a trajectory dependent fashion (Lipton et al., 2007).

Even in the absence of context units, i.e. with fully overlapping (the same) assemblies in competing sequences, our network can still solve a disambiguation task for sequences sharing two consecutive states in their trajectories (see the resonance phenomena in Fig.10F). While this phenomena allows the network to statistically solve sequence disambiguation for disambiguation windows of size 2, it does not generalize for longer sequential overlaps. One way to handle the problem in a more robust, consistent and transparent fashion is to use a mechanism that preserves the network’s dynamical history in a dynamical variable. In our future work we intend to add such mechanism to the network in the form of currents dependent on the z-traces that facilitate the longer maintenance of the information about past activations and thus support the disambiguation of sequences with more challenging overlaps.

### 3.4 Learning rule stability, competition and homeostasis

The stability of the learning dynamics of a firing rate network subject to associative learning tends to be accomplished by introducing weight dependent terms into weight updates (Van Rossum et al., 2000). This constrain is usually motivated and biologically interpreted as a homeostatic mechanism. Sequence learning models are not exempt from this necessity. One of the simpler approaches amounts to combining STDP with hetero-synaptic plasticity (Fiete et al., 2010). However, it is not straightforward how these two forces should be balanced. There are a plethora of models that rely on weight clipping with arbitrarily handpicked upper and lower limits (Mostafa and Indiveri, 2014; Veliz-Cuba et al., 2015; Murray et al., 2017). While this approach is analytically transparent, fine tuning between potentiation and depression is usually required. In a similar vein, Byrnes et al. (2011) introduced a combination of subtractive and multiplicative normalization as a mechanism of weight stabilization, which also has to be arbitrarily tuned. Verduzco-Flores et al. (2012) proposed a more complex approach that combines hetero-synaptic competition with a mechanism that limits both the total value of the weights and the total incoming current to a unit in order to achieve stability Pereira and Brunel (2018), on the other hand, resorted to a combination of synaptic normalization and multiplicative homoeostasis to avoid runaway excitation. While these two learning rules are able to prevent runaway instabilities and have varying degrees of biological plausibility, the number of parameters involved, and the complexity of the model are excessively high. As opposed to this complexity, the probabilistic nature of our BCPNN learning rule automatically accounts for weight competition during learning leading the network to a stable regime of sequential or attractor dynamics without requiring extra parameters or balancing different forces (as discussed more thoroughly by Tully et al. (2014)).

### 3.5 Limitations and further work

Although multiple studies of the cortical micro-circuitry have revealed distance dependent connectivity profiles (Xu et al., 2016; Jiang et al., 2015), we have ignored this design principle in our model. Previous spiking implementations of this model architecture have included to some degree both distance dependent effects in connectivity and distance dependent delays (Lundqvist et al., 2006; Tully et al., 2016; Fiebig and Lansner, 2017), which had impact on the network’s temporal dynamics. In our non-spiking network model the expected implications of such spatio-temporal diversity would be prolonged (temporally spread) attractor reactivation and transition processes. Still there should be no qualitative functional changes in the network’s behaviour as the key mechanisms would not be compromised (although see Spreizer et al. (2018) for a sequence production mechanism that arises itself from asymmetries in the spatial profile of connectivity). Due to the mesoscale nature of our model and interest in network phenomena, we obviously do not account for any dendritic related phenomena in sequence processing such as as the capacity of single neurons to work as sequence recognition devices through spatial effects (Branco et al., 2010) and the use of distal dendritic inputs to prime sequential activations (Hawkins and Ahmad, 2016).

In the presented work there are some phenomena that we have not systematically characterized in their generality. For example, in most simulations we exploited temporally homogeneous training protocols. To test the performance of our network under the conditions of varying pulse time, *T*_*p*_, and inter-pulse-interval, Δ*T*_*p*_, across patterns, we have ran preliminary tests and obtained promising results (data not shown). We intend to conduct a more comprehensive characterization of the network’s behaviour subject to highly variable training protocols (temporal pattern heterogeneity) in our future work.

## 4 Methods

### 4.1 Training and recall protocol

For our training protocol we created a time series **s**(*t*) to represent the input. **s**(*t*) encodes the information about the pulse time *T*_*p*_ and the inter-pulse interval IPI (Fig. 4B). We then performed off-line batch learning of the parameters using the integral formulation of the dynamic equations presented above (Eq. 6-7).

To avoid the ill-defined case for *p* = 0 we set the lower bound of *ϵ* = 10^-^7 for the argument of the logarithm. That is, if the value of *p* is less than *ϵ* we equate it to *ϵ*.

For training the two sequences with the overlapping representations we created the sequences in succession but separated among them by 1*s*. This ensured that the sequences in the training protocol were uncoupled from each other.

We say that pattern is active if the corresponding units are active for longer than *τ*_*s*_ (the smallest time constant in the system). The sequence is considered to be correctly recalled if by activating the first pattern all the others patterns in the sequence are subsequently activated in that given order. Given that for many possible tasks it suffices that the network state ends in the correct pattern or that only a part of the sequence is recalled correctly our success criteria is rather conservative.

### 4.2 Control and estimation of persistence time

In order to estimate the persistent time for a pattern *P* during recall we calculated the difference between the time *t*_1_ at which pattern *P* was activated and the time at which the next pattern was activated *t*_2_. *T*_*per*_ = *t*_2_ *- t*_1_.

As shown in Eq. 4, *T*_*per*_ time depends on both the weight and bias differences, Δ*w*_*next*_ = *w*_*self*_ – *w*_*next*_ and Δ*β* = *β*_*self*_ – *β*_*next*_ respectively and the adaptation gain *g*_*a*_. This offers flexibility in controlling the duration of patterns activations by adjusting the adaptation gain *g*_*a*_ as follows: 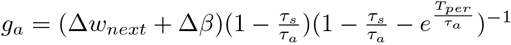. We use this adjustment to control *T*_*per*_ during recall in order to decouple the effects of training from the recall process.

### 4.3 Noise

Noise was included in our simulations as additive white noise with variance 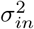 in the differential equation for the *s* variable. The current *s*, however, behaves almost as an Ornstein–Uhlenbeck (OU) process and therefore its standard deviation is given by 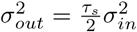. Based on this fact we characterized the effects of noise with the size of *σ*_*out*_ instead of *σ*_*in*_ The rational behind this choice is that *σ*_*out*_ will be closer to the standard deviation of the variable *s* in Eq. 1 and therefore comparable in magnitude to the value of currents in the network. It is important to say that thanks to the separation of times scales (*t*_*s*_ ≪ *t*_*a*_) the dynamics of *s* behaves mostly as an OU process and it is only the WTA dynamics around the transition points that induces deviations.

The incorporation of noise to the network makes the trajectories and, thereby, the recall process stochastic. To quantify the recall performance under noise (probability of successful recall at a given level of noise) we averaged the number of correct recalls in a given number of trials. The estimated probability of successful recall 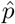 follows from a Bernoulli process and we can therefore quantify the uncertainty of our estimates with the Wald method to provide 95% confidence intervals (*N*_*trials*_ = 1000):

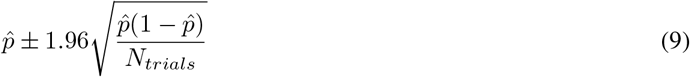

In order to systematically characterize how different parameters of our training protocol affect the sensitivity of the resulting network to noise, we estimated *σ*_50_ as the value of noise variance *σ* for which the probability of correctly recalling a given sequence is 0.5. Finding such *σ* is an instance of the Stochastic Root Finding Problem (Pasupathy, 2010). To estimate this we used the naive bisection algorithm for deterministic functions by using the averages as estimates of the actual values. We stopped the algorithm as soon as the success rate corresponding to our estimate of *σ*_50_ was contained in the Wald confidence interval given in Eq. 9. We find that our method was consistently able to find solutions to the root finding problem (see Fig. S 1 in the supplement).

To test for spread in the distribution of failure points we also calculated *σ*_75_ and *σ*_25_ (defined in an analogous manner to *σ*_50_) for the parameters under consideration. We found agreement in trend with our analysis (data not shown).

## 5 Aknowledgments

We thank Arvind Kumar for reading a draft of this work and providing valuable comments.

## Appendix

### A Complete treatment of the persistence time

To characterize the transition from pattern *m* to pattern *n* (standing for *P*_*m*_ and *P*_*n*_) in the units belonging to hypercolumn *i* we calculate the difference in their respective currents 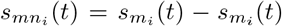. Where we have adopted the convention that *m*_*i*_ and *n*_*i*_ give the index of the unit belonging to pattern *m* and *n* in the hypercolumn *i* respectively. To obtain a solution for 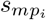 (*t*) we solve the resulting differential equation with the method of undetermined coefficients.

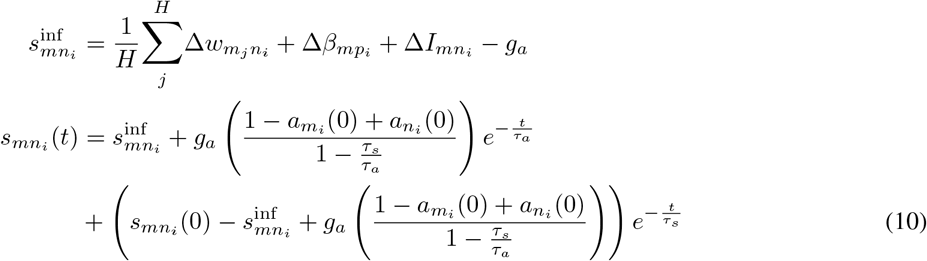

Where 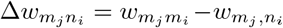 are the weights of the differential input coming to hypercolumn *i* from hypercolumn *j*, 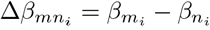 is the local (same hypercolumn) differential in intrinsic excitability and 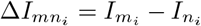 is the differential external input to the units belonging to *m* and *n* in the hypercolumn *i*.

When pattern *m* becomes active the units that belong to it start experiencing intrinsic adaptation through the terms 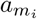 and in consequence 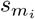 starts decreasing. It follows that the current 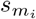 will become smaller than 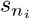 at some point in time and transition will occur. We denote such time as 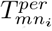 to emphasize that we are talking about transition from pattern *m* to *n* in hypercolumn *i*. Formally, this time can be found by setting 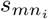, above equal to 0. If we disregard the short-term fluctuations of the term 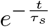 we obtain the following expression:

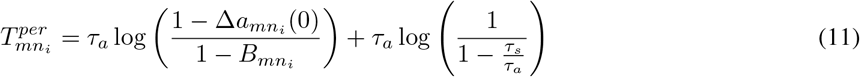

Where 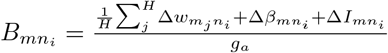 and 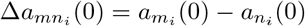. Note that the previous presence of adaptation in the unit of pattern *m*, 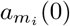, decreases the persistence time and previous presence of adaptation in the unit of pattern *n*, 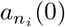, has the opposite effect.

In the case of multiple hypercolumns there is a value of 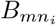 for every hypercolumn *i* determining how fast the transition happens at that hypercolumn. As a matter of fact the transition happens only if all the 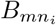 are less than 1. The transition is fast for 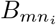 close to 0 and slow for 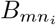 equal to 1 (modified by memory effects of the adaptation). These two effects combined will give the order in which the units of a pattern belonging to different hypercolumns undergo transition. However, the exact timings at which the transitions happen are modified after the first transition takes place; this is because the currents that the rest of the units of pattern *n* receive (the ones in the other hypercolumns) are modified as well. In general this will have the effect of accelerating the transition of the other units belonging pattern *n*. By taking this modifications into account we can derive conditions for the modification of *Tper* in the remaining hypercolumns after a transition in hypercolumn *k* has happened (up to time differences in the order of *τ*_*s*_ due to membrane capacitance effects):

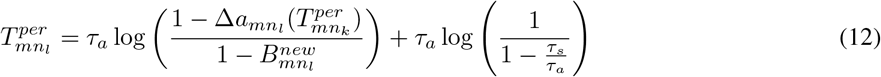

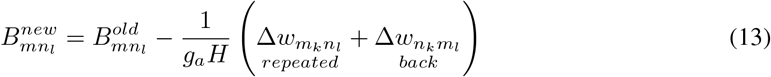

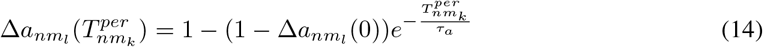

The 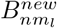 term is now reduced by the lost self-excitatory current from unit *m*_*k*_, 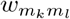 (we also subtract the lost of the feed-forward current 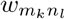) This reduction is reflected in the subtraction of the term 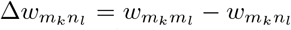. The now activated unit *n*_*k*_ induces a backward current: 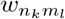. Also there is a recurrent current helping to fix the *m*_*l*_ unit coming from hypercolumn *k*, 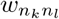. These contributions are reflected in the addition of the terms 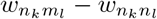 to the expression above which we write with a minus sign as: 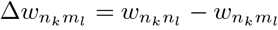. The overall effect of these new currents (mainly coming from the backwards negative current 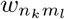) is to reduce the value of 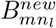 with respect to 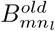 thus effectively hastening the transition. Moreover, as time passes, the adaptation current tends to become larger in the units that are activated and smaller in the units that are not which also contributes to speed up the transition. This effect is reflected in the quantity 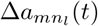 becoming closer to 1. We can use this effect iteratively to calculate the values of 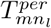 for every hypercolumn using the formula above recursively.

To derive conditions for synchronous transition we notice that if the term inside the logarithm becomes less than 1 it means that the quantity becomes negative implying instantaneous transition. This is accomplished when the following condition is satisfied:

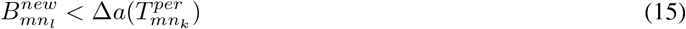

As long as there is a hypercolumn for which this value is satisfied the transition takes place there. This in turn, means that the values of *B* have to be updated again (making them smaller) rendering a transition in the other hypercolumns more likely. This creates a cascade effect where the latter transitions happen overwhelmingly faster than the first ones.

Please note that while this provides us with transition times for all the hypercolumns between two patterns, it does not guarantee that the aforementioned transitions will be the ones that happen. It is still possible that other values of 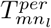 are smaller and those are the transitions that in fact occur.

## Supplementary figures

**Figure S1:**
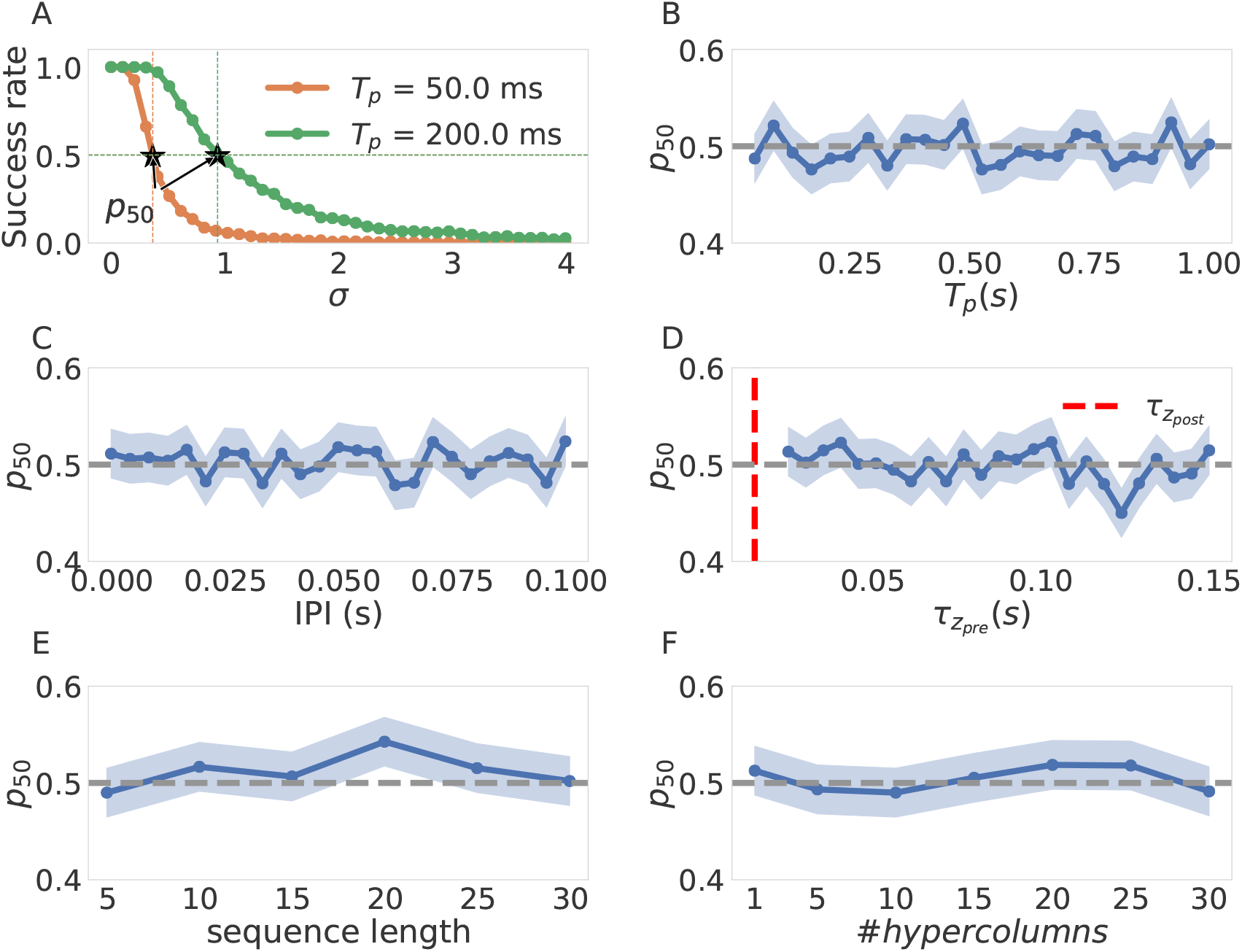
Calibration of *σ*_50_estimation. (A) two success rate vs noise profiles for *T*_*p*_= 50 *ms* and *T*_*p*_= 200 *ms*. The values of *p*_50_are annotated for reference. (B-F) We show the values of *p*_50_obtained after running the algorithm in Fig. 8. For every value we see that the values of the found roots (*p*_50_, blue lines) was within confidence bounds (here blue shaded) of the expected value (0.5, horizontal lien in gray).

